# Online Bayesian Optimization of Nerve Stimulation

**DOI:** 10.1101/2023.08.30.555315

**Authors:** Lorenz Wernisch, Tristan Edwards, Antonin Berthon, Olivier Tessier-Lariviere, Elvijs Sarkans, Myrta Stoukidi, Pascal Fortier-Poisson, Max Pinkney, Michael Thornton, Catherine Hanley, Susannah Lee, Joel Jennings, Ben Appleton, Phillip Garsed, Bret Patterson, Will Buttinger, Samuel Gonshaw, Matjaž Jakopec, Sudhakaran Shunmugam, Jorin Mamen, Aleksi Tukiainen, Guillaume Lajoie, Oliver Armitage, Emil Hewage

**Affiliations:** BIOS Health Ltd; Université de Montéal and Mila-Quebec AI Institute

## Abstract

**Objective:** In bioelectronic medicine, neuromodulation therapies induce neural signals to the brain or organs modifying their function. Stimulation devices, capable of triggering exogenous neural signals using electrical wave forms, require a complex and multi-dimensional parameter space in order to control such wave forms. Determining the best combination of parameters (wave form optimization, or dosing) for treating a particular patient’s illness is therefore challenging. Comprehensive parameter searching for an optimal stimulation effect is often infeasible in a clinical setting, due to the size of the parameter space. Restricting this space, however, may lead to sub-optimal therapeutic results, reduced responder rates, and adverse effects.

**Approach:** As an alternative to a full parameter search, we present a flexible machine learning, data acquisition and processing framework for optimizing neural stimulation parameters requiring as few steps as possible using Bayesian optimization. Such optimization builds a model of the neural and physiological responses to stimulations enabling it to optimize stimulation parameters and to provide estimates of the accuracy of the response model. The vagus nerve innervates, among other thoracic and visceral organs, the heart, thus controlling heart rate and is therefore ideal for demonstrating the effectiveness of our approach. *Main results.* The efficacy of our optimization approach was first evaluated on simulated neural responses, then applied to vagus nerve stimulation intraoperatively in porcine subjects. Optimization converged quickly on parameters achieving target heart rates and optimizing neural B-fibre activations despite high intersubject variability.

**Significance:** An optimized stimulation waveform was achieved in real time with far fewer stimulations than required by alternative optimization strategies, thus minimizing exposure to side effects. Uncertainty estimates helped avoiding stimulations outside a safe range. Our approach shows that a complex set of neural stimulation parameters can be optimized in real-time for a patient to achieve a personalized precision dosing.

## 1 Introduction

Bioelectronic medicine applies neuromodulation to modify neural activity and physiological function, providing a novel approach to treating a range of neurological and physiological diseases. In contrast to exploiting endogenous (or spontaneous) neural activity, stimulation devices are able to trigger exogenous neural signals. For the stimulation of nerves, exogenous neural signals take the form of evoked compound action potentials (eCAP), the result of a depolarization of a large number of fibres in the nerve triggered by an electrical stimulus (examples of eCAPs are shown in Fig. 4). The type and number of fibres recruited depend on the wave form and dosing of the stimulus in a complex way. Through such signals, therapies utilizing stimulations can alter the function of organs, offering promising new avenues for medical treatment (Cracchiolo et al., 2021). The exogenous signal is typically delivered by an implantable pulse generator (IPG) via electrodes placed with nerve tissue.

The autonomic nervous system (ANS) is essential for maintaining homeostasis throughout the body and is implicated in the pathogenesis of a variety of disorders. The vagus nerve (VN) is a key target for neuromodulation of the ANS, as it innervates several thoracic and visceral organs, including the heart, lungs and gut. Accessing the VN through surgery is relatively straightforward (Reid, 1990) and vagus nerve stimulation (VNS) has been suggested for the management of neurological diseases such as drug resistant epilepsy (Labiner and Ahern, 2007a), depression (Nemeroff et al., 2006), as well as respiratory (Chang et al., 2015; Carr and Undem, 2003), inflammatory (Pavlov and Tracey, 2012) and cognitive conditions (Meyers et al., 2018; Martin et al., 2022), among others.

Heart failure (HF) has been a primary target for VNS therapy in several preclinical and clinical studies. A specific change in heart rate (HR) immediately after VNS is the basis of most VNS therapies for heart failure suggested to date: cyclic application of stimuli that achieve mild bradycardia have shown long term therapeutic effects in preclinical studies (Dusi and De Ferrari, 2021). Despite promising results in early human trials, phase II and phase III controlled trials did not confirm efficacy (Dusi and De Ferrari, 2021). The stimulation doses that could be reached in clinical trials were markedly lower compared with those applied in preclinical studies due to side effects reported by the patients, such as cough and voice alteration. In particular, based on earlier studies (Ardell et al., 2017a; Nearing et al., 2020), stimulations that achieve a balance between tachy- and bradycardia for an overall net effect of zero change in heart rate are the focus of the most recent clinical trial (Konstam et al., 2019), which aims to address some of the shortcomings of earlier clinical trials.

While the potential of VNS is widely recognized, important challenges remain in developing effective stimulation protocols. Notably, there is substantial patient-to-patient variation in the distribution and arrangement of fibres in the VN (Upadhye et al., 2021; Jayaprakash et al., 2022), as well as in the coupling of these fibre groups and stimulation electrodes (Qiao, Stieglitz, and Yoshida, 2016). This necessitates a personalized optimization process. The optimization of VNS for HF therapies is primarily based on mitigating potential off-target effects prior to considering the efficacy of the resulting VNS setting (Labiner and Ahern, 2007b; Nicolai et al., 2020). Besides patient to patient variation in the effect of VNS parameters, changes within patients over time have been observed as well (for example, over a ten week titration period in Nearing et al. (2020)). Extensive adjustments over a large number of stimulation parameters are time consuming and often induce discomfort or even more severe side effects. The development of rapid and flexible optimization procedures which can quickly and efficiently identify stimulation parameters with a specific response is therefore of great interest.

Personalising neural stimulation settings is a complex task. BIOS’s neural stimulation optimisation system is comprised proprietary neural interface which collects high quality longitudinal neural and physiological data, a library of neural biomarkers, cloud infrastructure designed specifically for time syncing neural and physiological signals, and optimisation algorithms to efficiently find personalised settings in narrow surgical windows. While all of those pieces together enable stimulation optimisation, the focus of this paper will be a specific optimisation method—Bayesian optimization (BO). Here we label our specific version of Bayesian optimisation applied to neural stimulation as Online Bayesian Optimisation of Evoked Signals (OBOES).

Bayesian optimization is a computational technique that seeks to identify optimal parameters with a minimum number of exploratory steps. BO algorithms have recently emerged as promising optimization tools for stimulations of the nervous system in several studies: for optimizing epidural spinal cord stimulation in rats (Desautels et al., 2015), for intracortical stimulation in monkeys (Laferriere et al., 2020), for distal limb movement in monkeys (Losanno et al., 2021), and for compound action potentials (Mao et al., 2022). The key elements of our OBOES algorithm are as follows. At the outset of an optimization procedure for a patient their VNS response function is unknown, necessitating some form of exploration. At the same time, steps need to be taken towards optimizing the function. Thus, a balance must be struck between exploration and optimization. Finally, a statistical assessment of the extent of exploration and of the confidence range on the discovered optimum is desirable. Knowledge of the probability that responses will likely fall outside a safety or comfort range helps managing the range of acceptable dosing.

We develop and tested OBOES by optimizing responses to stimulations controlling heart rate and eliciting optimized activations of certain nerve fibre types. To develop, compare and evaluate optimization approaches *offline* (or *in silico*), we developed realistic simulation models of typical neural responses to VNS using experimental data. These simulation models are data-driven with minimal modelling assumptions about any underlying mechanisms, thus providing a realistic evaluation platform for assessing the effectiveness of optimization approaches for various VNS scenarios. The effectiveness of a neural stimulation optimisation using a Bayesian method approach in an *online* (or *in vivo*) setting was explored in a series of VNS experiments conducted on anaesthetized swine, the preferred animal model for investigating the function of the human vagus nerve. Swine and human vagus nerves and overall anatomy are highly similar—making porcine models suitable for translation.

In addition to the optimization of changes in HR, we performed optimization of neural biomarkers, such as the activation of specific fibre types. To this end, we recorded exogenous nerve signals induced by VNS, in the form of eCAPs. It has been widely hypothesized that B-fibre activation is associated with bradycardia (Qing et al., 2018). Fibre activation in the exogenous signal can be optimized with minimal physiological effects using stimulations with low frequency and short duration. Optimizing fibre activation instead of directly targeting a change in heart rate is therefore less stressful for the subject and less time consuming. As an example of the applicability of OBOES to optimize specific neural signals in an online experiment, we maximized B-fibre activation while minimizing stimulation current.

Deployment of OBOES was enabled by a R&D system comprising the implantation of nerve cuffs, data acquisition of neural and physiological data, and data and algorithm management in the cloud.

In summary, we have demonstrated the practical applicability of a machine learning optimization algorithm for VNS in a series of preclinical trials. This system achieves, in real time, target changes in heart rate or maximization of B-fibre activation while minimizing stimulation current. Consequently, the system facilitates the optimization of multi-dimensional VNS parameters, thus enabling a personalized precision dosing with a minimum number of stimulations causing side effects.

## 2 Methods

We describe the surgical setup and technical infrastructure for recording neurograms of evoked compound action potentials (eCAPs) in the VN and heart rate changes with VNS. To assess the performance of our optimization approach on neural data we built a series of simulators. Since the ”true” response function is known for simulators, optimization algorithms can be compared and assessed and the efficiency of an optimization can be evaluated *offline*. Simulators for eCAPs were constructed based on VNS responses recorded from our subjects, termed the *Porcine Dataset*. A further set of simulators is based on data available in the literature, termed the *Rodent Dataset*. Finally, we performed *online* optimizations of heart rates and neural fibre activations intra-surgically and describe a statistical method for assessing the uncertainty estimates of OBOES when the true response function is unknown.

### 2.1 Anaesthesia and surgical preparation

Experiments were conducted on five female Landrace pigs (labelled A7 to A11) under the approval of the local Institute of Animal Care and Use Committee, and approved in accordance with the animal use policy of BIOS Health Ltd. All animals, weighing 43.2*±*3.5 kg, were sedated using an intramuscular injection of Ketamine (25 mg kg*^−^*^1^), Atropine (0.04 mg kg*^−^*^1^) and Acepromazine (1.1 mg kg*^−^*^1^), then intubated and anesthetized using Propofol (3 mL slow intravenous bolus for induction and 0.4 mg kg*^−^*^1^ min*^−^*^1^ for maintenance). The depth of anaesthesia was monitored and adjusted based on the occipital reflex, jaw tone and haemodynamic indices. A mechanical respirator was available, although animals were left to breathe spontaneously unless end tidal carbon dioxide concentration exceeded 70 mmHg. A two-lead electrocardiography (ECG) was used in a Lead I configuration. Data were digitized at a rate of 500 Hz using the Axon Digidata 1550B plus HumSilencer (Molecular Devices, CA, USA).

Animals were positioned supine, with both forelimbs and head extended to expose the ventral aspect of the neck. A ten centimeter incision was marked two centimetres right of the midline. The nodose ganglion marks the most cranial point of our interaction with the vagus. Eight centimetres of the nerve was stripped and cleaned caudally of the nodose to fit three cuffs.

The surgical procedures each ran for eight hours at which point the animals were euthanized with an intravenous administration of T-61 (0.3 mL kg*^−^*^1^).

### 2.2 Electrodes and equipment

The multi-contact cuffs, built by Microprobes (NC, USA), each contained eight electrodes arranged in longitudinal bipolar pairs (see Fig. 1). All neural recordings and stimulation data presented in this paper were performed via the maximally spaced bipolar pair (electrodes 1 and 8 in Fig. 1), with a centre-to-centre separation of 6.2 mm. The cuff arrangement gave an average inter-cuff distance of 24.5*±*15.8 mm between the stimulation cuff and the most cranial recording cuff across the cohort.

**Figure 1:**
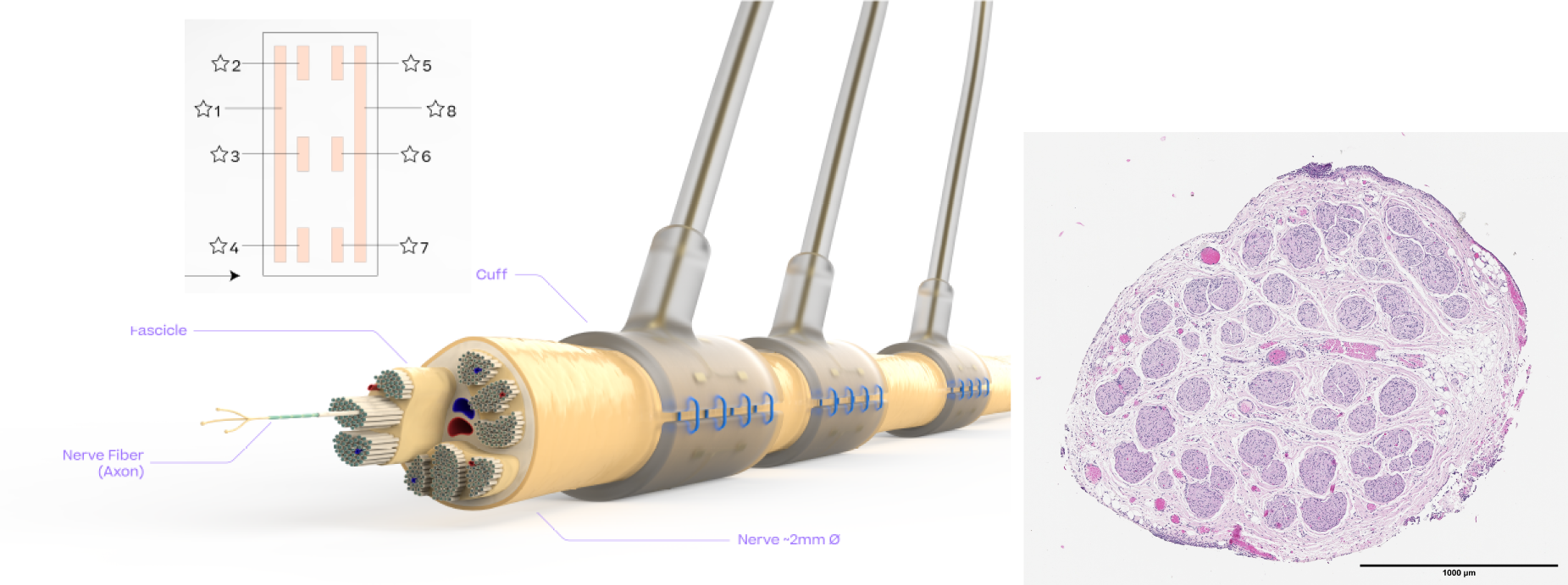
Illustration and histology of the cross section of the vagus nerve showing cuff placement and electrode layout. The multi-fascicle nature of the vagus nerve poses a challenge for the targeted activation of a specific fibre or fascicle within the bundle over other fibres, given the size and location of electrode contacts relative to the fibre positioning.

All cuffs were connected to a custom Data Acquisition System engineered in-house. Briefly, a RHD Amplifier and RHS Stimulation and Amplifier chips (Intan Technologies) are controlled through custom HDL on an Artix 7 (Xlinix, CA, USA) FPGA sampling up to 16 channels at 30kHz. The system is capable of applying stimulation pulses with varying current (10–2500 µA), frequency (1–1500 Hz), individual pulse width (1–1000 µs), and train duration (1–10 s). Biphasic symmetric constant-current pulses were delivered through a single channel in reference to a ground lead positioned proximal to the stimulation site. A spare pair of record channels were used as an additional ECG recording to ease synchronisation of the neural and physiological data and to obtain heart rate data during online optimization.

### 2.3 Research and development system architecture

We developed a software suite to accelerate our R&D process with the goal of coupling data generation with cloud-based data management and processing resources to support machine learning research in neuroscience. A schematic of the platform is shown in Fig. 3.

The data acquisition components of the system reside in the operating theatre: the Data Acquisition System (Neural Interface) and User Interface (UI, NeuroTool). The Data Acquisition System performs the VNS and VN signal recording, uploads data to the cloud services and has an onboard GPU for edge ML inference. Stimulation control models are deployed here for applications where rapid inference and data security are paramount. Data can be temporarily streamed to the UI for validation and checking of impedances. All stimulation parameters can be programmed, adjusted and triggered by an onsite electrophysiologist.

Data are uploaded to a Data Ingestion Service in batches. From here, the neural responses of interest can be immediately visualized and examined on *Data Explorer* or on *OBOES Explorer* (see Fig.3; snapshots of the explorer screens are shown in the supplementary material). Monitoring of responses informs fibre activation thresholds to guide other stages in the protocol. Data are stored on a remote server and can be downloaded for post surgical analysis and algorithm development.

### 2.4 Rodent Dataset

For the development of the OBOES framework, we devised a simulation platform for testing, refining and evaluating optimization approaches on simulated data. Such simulations should show characteristics as close as possible to realistic neural or physiological responses to stimulations. We utilized data on neurograms of evoked compound action potentials (eCAP) by Ward et al. (2015), available through the NIH SPARC programme (Ward et al., 2021). The dataset provides the responses to a grid of VNS parameter values varying over seven pulse currents and four pulse widths. Each stimulation was applied cathode-first with an alternating monophasic constant current. Twelve rodents received the complete sweep of 28 VNS parameter combinations at a fixed pulse frequency of 10 Hz. Here we refer to the rodents as subjects P1 to P12. The maximum eCAP values provided in the study form the basis of our simulator construction.

### 2.5 Porcine Dataset

We extended the simulation platform using data from our own experiments. Exogenous signals in the form of evoked compound action potential (eCAP) responses to stimulation were collected for four porcine subjects A7 to A11. The pulse width was set to either 130 µs, 260 µs, or 500 µs and the pulse train duration to five seconds. A range of frequencies (2, 5, 10, 15, 20 Hz) and currents (50–2500 µA, in increments of 100 µA) were applied. We found an alternating, monophasic, constant pulse waveform to be the most suitable to elicit clean eCAP responses without amplifier artefacts.

Evoked neural and cardiovascular responses to VNS were recorded and vitals were allowed to return to baseline before starting the next cycle (with a minimum of 30 seconds between each cycle). Charge limits were set at the point where stimuli caused noticeable strong side effects, such as coughs or extended bradypnea. Neurograms were processed as described next.

### 2.6 Processing of neural data

Neural responses to individual stimulation pulses were identical and only the eCAP response to the first pulse in a pulse train was used for simulation. Any trends were removed by robust polynomial regression (Seabold and Perktold, 2010). This is followed by convolution smoothing using a Kaiser window of increasing size, in order to retain fine details at the beginning of the signal for faster, less dispersed eCAPs, while providing enough smoothing towards the end of the signal for slower, more dispersed eCAPs. As an example, the evolution of the fibre engagement for two subjects, recorded from the most cranial cuff, are displayed in Fig. 4. Additional figures of eCAPs for all subjects can be found in the supplementary material.

Activations of three fibre types are visible in the neurograms of Fig. 4, denoted A*β*-, A*γ*-, B-fibres followed by a laryngeal muscle artefact. The muscle artefact was validated through caudal vagotomy as in (Nicolai et al., 2020). We noticed that the artefact appears instantaneously on both cuffs without any conduction delay in contrast to the other eCAPs. Fibre activation thresholds differ from subject to subject. This is likely due to differences in bioelectronic coupling at the cuff-nerve interface and anatomical variation in fibre distribution (Losanno et al., 2021).

For optimizing B-fibre activation, activation was calculated from neurograms by identifying the B-fibre maximum and minimum peak and taking their difference.

### 2.7 Gaussian Processes

Gaussian processes (GP) (Rasmussen, 2006) provide a flexible, nonlinear approximation of an unknown response function. Such function, seen as a black box, provides response values to given inputs. With only a finite number of inputs and corresponding response values, limited information about the underlying function is available, and the approximation might not be very accurate. If subsequently more data become available, it is possible to integrate them in a principled probabilistic fashion to improve the GP approximation. Here we use GP approximations for two purposes: a) constructing simulation models for method development and testing, described in section 2.8, and b) serving as a surrogate function in Bayesian optimization, as outlined in section 2.9.

Technical details of how a GP approximation is set up can be found in the supplementary materials. Essentially, a GP provides an approximation to a response function *g* that maps a *d*-dimensional input *x ∈* ℝ*^d^* to to a scalar response *y*, *y* = *g*(*x*). A key feature of a GP is the ability to provide an estimate of the precision of its approximation in the form of uncertainty regions above and below its approximation. The accuracy of the approximation improves with an increasing number of data points of input-response pairs. The GP is represented by a mean function *m*(*x*) for any input *x* and an uncertainty estimate *ν*(*x*) at this point. The smaller *ν*(*x*) the more accurate the approximation. In this study GP approximations are shown as surfaces (of mean response) with upper and lower confidence bounds indicated as transparent surfaces when the uncertainty around the GP mean response is of interest. They consist of the surfaces for 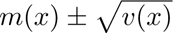 enclosing the central 68.2% quantile of a normal distribution. For examples showing of all three surfaces see Fig. 8.

Fitting a GP to data (*x, y*) means choosing suitable kernel parameters that represent the variance of the signal, the multi dimensional length scales (smoothness), and the noise variance. Parameters can either be set manually or estimated by maximum a posteriori probability (MAP) with or without a prior such as a Gamma distribution (some details on priors used in this study can be found in the supplementary material).

### 2.8 Simulators for the Rodent and Porcine Dataset

In order to develop and assess optimization approaches offline, we developed simulators based on the Rodent as well as on our Porcine Dataset. For the Rodent Dataset we constructed GP simulators for maximal eCAP activation for each of the 12 subjects P1 to P12. A GP approximation of maximal eCAP action in subject P10, for example, is shown in Fig. 6c. The original data provide 28 data points (shown in blue in Fig. 6c). A GP approximation provides an output for any arbitrary input within a given search space. For more realism the simulator adds a random value at a noise to signal ratio (NSR) of 0.2 to the output at each request as observed in the dataset. Results for an NSR of 0.1 to 0.4 look qualitatively very similar and are not shown.

For the Porcine Dataset of subjects A7 to A10 we constructed GP simulators that include the time dimension as well to represent the neural signal as time series, that is, a three dimensional input point of current, width, and time is mapped to the corresponding electrode voltage value at that time (for an example see Fig. 5). For further details of GP parameter settings see the supplementary material. For an input of stimulation parameters this type of simulator returns a time-series vector, a neurogram of eCAPs. In order to obtain a scalar response representing B-fibre activation, the activation value was computed as difference of the maximum in the first half to the minimum in the second half of the B-fibre range defined as 2.47–4.3 ms, 2.3–4.97 ms, 2.97–5.97 ms, and 3.97–6.3 ms for A7 to A10.

### 2.9 Bayesian Optimization

Bayesian optimization (Archetti and Candelieri, 2019) is a popular choice for the optimization of objectives over unknown functions. BO is a sequential search strategy optimizing the input to an unknown response function balancing optimization with exploration (Mockus (2012), for an accessible introduction see also BorealisAI (2020)). Typically the response function to be optimized over is unknown. However, it can be probed at any arbitrary input point (possibly with certain boundaries). In the context of a BO the mechanism producing responses to inputs is referred to as the *oracle* or *black box* function. In what follows, we will use BO in two contexts: when the oracle is provided *offline* by a simulator (a GP fitted on a full data set), and when the oracle is provided *online* by a real-time measurement of the physiological or neural response following VNS in an alive subject. For details, see sections 2.11 and 2.12, respectively.

Since the optimizer has no access to the true response function and can only query it at a few input points through the oracles, it builds a *surrogate function* which it maintains and improves internally to guide its search. Sequential querying of the oracle is used to improve the surrogate function. The surrogate function therefore needs to be flexible enough to adapt to new data. It should also provide some estimation of uncertainty in order to balance optimization with exploration. This is required in order to detect regions of high uncertainty that might require more data to narrow the uncertainty gap.

### 2.10 Bayesian Optimization with Gaussian processes

GPs are a popular choice for representing surrogate functions in BOs. As discussed in section 2.7, GPs are probabilistic models that are able to capture functional relationships between multidimensional inputs and a nonlinear responses. Starting from a typically flat representation of the function before any data are available, a GP provides an increasingly accurate estimation of the unknown response function by incorporating an growing number of data points.

A BO run consists of a series of requests in the form of queries to the oracle. To find the next query input to forward to the oracle, a BO maintains an *acquisition function* in addition to its surrogate function. The next query input is an input that optimizes the acquisition function. The acquisition function needs to support finding an optimum of the (unknown) objective function based on the BO’s surrogate function. However, it also needs to support exploration of new regions for an improved estimation of the surrogate function. To achieve these complementary aims an acquisition function is typically some combination of the mean of the surrogate function with its uncertainty. Once the acquisition function is optimized and the corresponding input is forwarded as query to the oracle, the oracle returns the response to that input. The acquisition function is designed so that this response either provides an improved optimum, or reduces the uncertainty about the response function in an under-explored region. The new data point contributes to an improved estimation of the objective function by the surrogate function.

The upper (lower) confidence bound UCB (LCB) is a popular acquisition function if the goal is to maximize (minimize) the objective. As in section 2.7 a GP is given in form of its mean function *m*(*x*) and the uncertainty variance *v*(*x*) or standard deviation 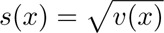 at input *x*. LCB and UCB are then defined as lcb(*x*) = *m*(*x*)*−λs*(*x*) and ucb(*x*) = *m*(*x*)+*λs*(*x*) (*λ* a scaling factor that trades off exploration with exploitation; we call it the *confidence bound factor*).

Since we often not necessarily aim to maximize or minimize a response, but to get close to a specific set point, we propose a novel acquisition function. The *targeted confidence bound* method or TCB, is a variation of the UCB and LCB acquisition function and defined as tcb(*x*) = *|m*(*x*) *− t| − λs*(*x*). An optimal query point is found as *x^∗^* = argmin*_x_*tcb(*x*). A justification for the use of this acquisition function to find an optimal point achieving a target set point is provided in the supplementary material.

### 2.11 Offline Bayesian optimization: assessment using simulations

Before deploying OBOES in a live experiment, we assessed the feasibility, practicability, and the convergence rate to be expected for typical neural response functions. We used the simulators from section 2.8 to provide challenging objective functions from realistic neural data sets. With a simulator, we have access to the true objective function and can therefore provide the ’ground truth’ optimum. This optimum can be used as a yardstick to assess how quickly an optimization algorithm converges to it in terms of numbers of required queries.

We expect the number of queries required by an efficient optimization algorithm for reaching an optimum to be small. Using a simulator, an evaluation of BO in terms of the required number of iterations was done as follows. First we obtained the true optimal value from the simulator. Then, starting with two initial queries at the extrema of lowest and highest values (for all input dimensions) within the search space, we tracked the progress of BO iterations from query to query. At each iteration we obtained the BO’s suggestion for the next query point (which optimizes its acquisition function) as input to the oracle in the next iteration. For the purpose of assessment, however, we also obtained the BO’s best current guess of an input that might elicit an optimal response from the oracle (which optimizes its surrogate function). Since the best current guess is obtained by optimizing the surrogate function, the estimated optimal input typically differs from the query input, which is obtained by optimizing the acquisition function. For the assessment, the algorithm’s estimated optimal input is evaluated by the simulator. The value returned for that input is compared to the true optimum (which was obtained from the simulator in the beginning). The relative error is the absolute difference between guessed and true optimum divided by the true optimum.

The most straightforward alternative to a BO search is a *grid search* with a fixed number of input points laid out in a grid. Notice that optimization using a grid requires a large amount of oracle evaluations before an optimum can be called, since all grid points need to be evaluated. The relative error of this optimum serves as a threshold any competitive optimization method should be able to undercut in fewer iterations than the size of the grid. A *random search* evaluates random input points sequentially through the oracle and keeps track of the best result along the sequence. For examples of a comparison of all three search strategies see Fig. 7. We expect that the relative error of a BO is smaller than that of a random search for the same number of iterations.

### 2.12 Online Bayesian optimization: modulating heart rate change and B-fibre activation

Two types of online experiments were conducted using OBOES. The first experiment aims to optimize VNS to meet a target modulated heart rate change. The second aims to optimize a specific vagal fiber recruitment, as measured by recorded eCAPS.

Online experiments for heart rate were conducted in subjects A9, A10, A11 and for B-fibre optimization in subject A12. A safe search space for pulse frequency and current was established based on previous stimulation experiments and the judgment of an onsite electrophysiologist. We initialized the BO algorithm with two fixed stimulation settings: one at lowest frequency and current setting, one at the highest within the search space. The optimization cycle proceeds as shown in Fig. 2. To achieve a target heart rate, a target value for ΔHR was chosen: *−*5 bpm for A9 and A11, and *−*10 bpm for A10. After initialisation, the BO model suggests a query input, consisting of a current (I/µA) and a frequency (f/Hz). A stimulation with these query settings is applied. The cardiovascular and vagus response for this stimulation are uploaded via the Data Ingestion Service serving OBOES Explorer (Fig. 3). The heart rates on either side of the stimuli are estimated from ECG data (for details see supplementary material) and their difference ΔHR returned to the BO algorithm as response to the query.

**Figure 2:**
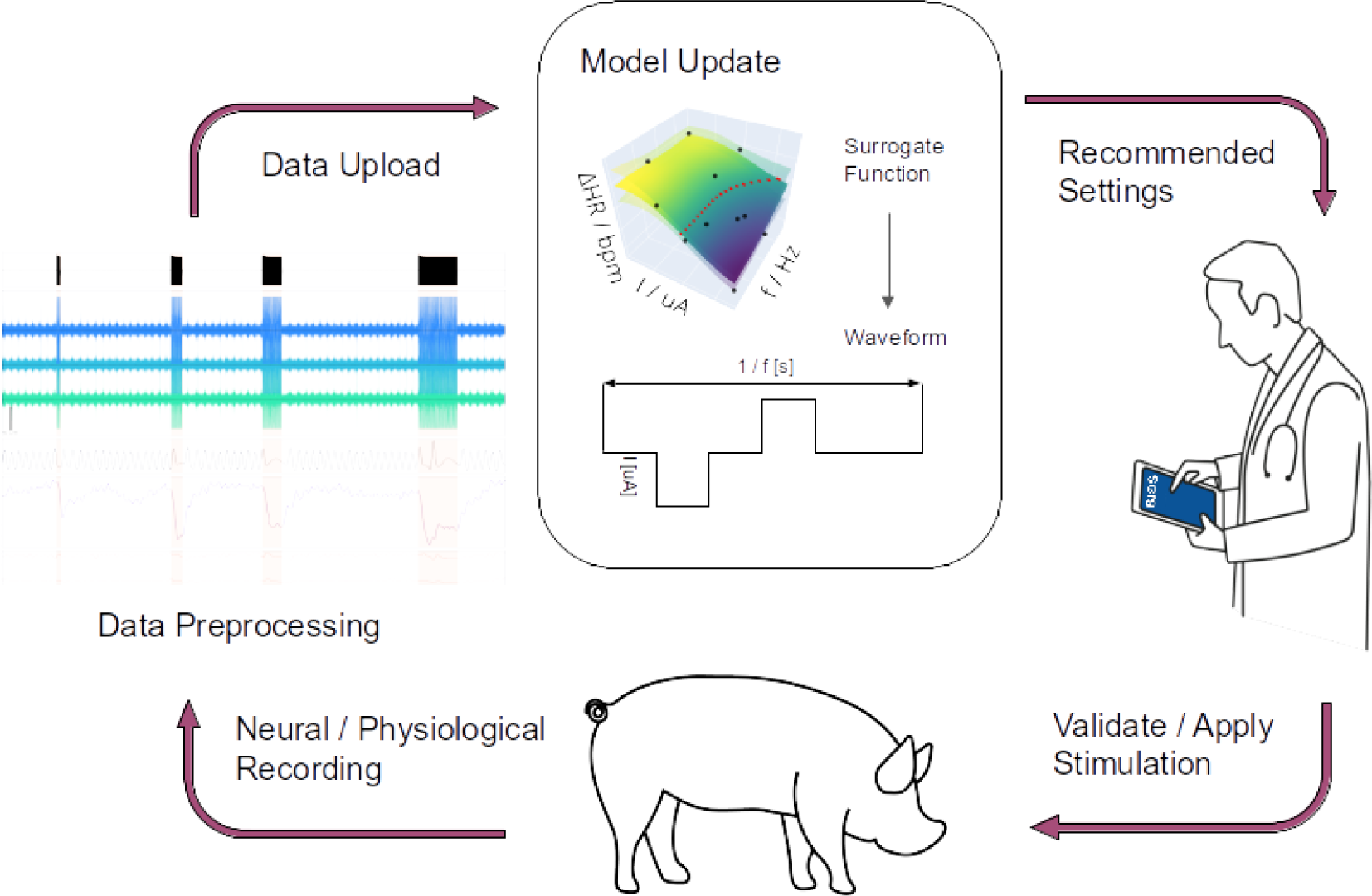
Overview of OBOES framework. Device parameters (a query) for the next stimulation event are suggested by BO. If the operator deems the stimulation parameters to be safe considering the current state of the subject, they apply the neural stimulus. Neuronal as well as physiological responses are recorded. The optimizer integrates the new response in an improved surrogate function: a model representation of the true response function. The BO algorithm then provides a new query that will result in a response that either contributes to an improved accuracy of the surrogate model function or is closer to the target value. Here OBOES is illustrated for a target value of a specific (red dots) change in heart rate (ΔHR) over pulse frequency (f) and current (I).

**Figure 3:**
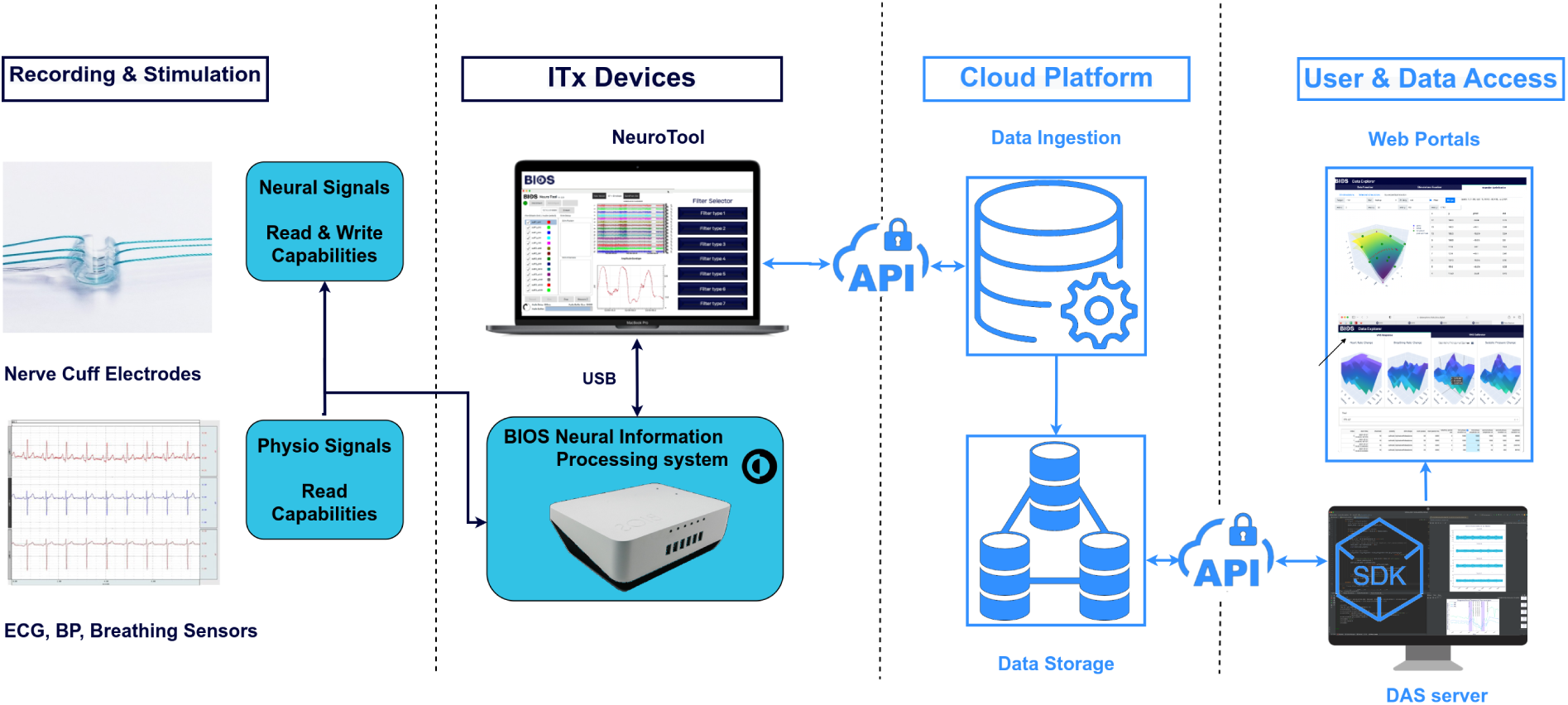
High-level overview of the ITx Platform. Neural and physiological data are collected using proprietary components deployed in the operating theatre and uploaded to the cloud environment, where it is accessed via various portals. *NeuroTool:* system for delivering stimulation protocols, monitoring vitals, neural recordings and in the operating theatre, *Web Portals:* OBOES Explorer for running and monitoring the optimization, Data Explorer for analysing and displaying raw and processed data.

Similarly, OBOES was set up for maximization of B-fibre activation as recorded by cuff electrodes. Neurograms were recorded during stimulation and processed as described in section 2.6. B-fibre activation was defined as the difference between maximum and minimum value of the recording in the range of 3.5–5.2 ms. In order to mitigate side effects due to strong stimulation currents, the objective function was defined as the ratio of B-fibre activation (in mV) and stimulation current (in µA). This objective function encourages high levels of B-fibre activation and simultaneously low levels of stimulation current.

There is no immediate ground truth for assessing an online BO. However, a targeted BO aims for a specific target response (for example see red points in Fig. 6b). Instead of a single input that leads to a maximum or minimum response the BO typically provides a range of inputs along the iso-response curve with responses close to the target value. We can therefore choose from a range of input values to assess the BO. Ideally we would like to demonstrate the accuracy of the BO’s surrogate function by establishing that the true response value falls within the uncertainty range of one standard deviation above and below the mean of the surrogate function GP of the BO (see section 2.7). However, the experimental response to VNS is affected by noise, and a single response value is not necessarily representative of the true response. We therefore acquired three responses *r*_1_(*x*)*, r*_2_(*x*)*, r*_3_(*x*) for the same input *x* and used them to estimate the true response value *ρ*(*x*) at that input by Bayesian inference. Instead of a single value, such inference results in a probability distribution for *ρ*(*x*) (for details see supplementary material). We consider the accuracy of the surrogate function as sufficient if 95% of this probability mass lies within the BO’s uncertainty range. This uncertainty range is then shown to be a reliable measure of how close the BO’s surrogate function is to the real response, at least around the iso-response curve. We repeated this calculation for three different input points along this curve. For an illustration of this approach and an example of the posterior distribution see Fig. 9b.

For maximizing the B-fibre response over current, we obtain a single optimal input instead of an iso-response curve of possible inputs. So instead of testing the accuracy of the surrogate function and the uncertainty estimate at three different inputs along an iso-response curve, we just tested the accuracy of the surrogate function at the optimal input.

## 3 Results

In order to assess OBOES offline, realistic simulators were constructed from neural data as reported in the following section. Subsequently, the results of evaluating the approach offline are discussed, followed by sections detailing the results from applying OBOES online in an intra-operative setting with alive animals, controlling heart rate and B-fibre activation.

### 3.1 eCAPs simulators for Porcine Dataset

For assessing the optimization of neural fibre activations through VNS we built simulators emulating eCAP responses. eCAPs for four different subjects A7 to A10 were obtained for this study. Fig. 4 shows eCAPs following VNS for subject A8 and A10 for pulse widths of 260 µs and 500 µs. Fig. 5 shows the result of fitting a GP to a three dimensional input space of current, width, and time to provide simulated voltage reading at this point for A10. Fig. 5a and 5b show the fit to the two different levels of pulse widths available in the data set. However, notice that the simulator can provide eCAPs for arbitrary inputs: Fig. 5c shows a simulated eCAP response for pulse width of 310 µs and current of 2.13 mA, neither of which was covered by the original data. Figures of eCAP simulators for A7, A8, and A9 are provided in the supplementary material.

**Figure 4:**
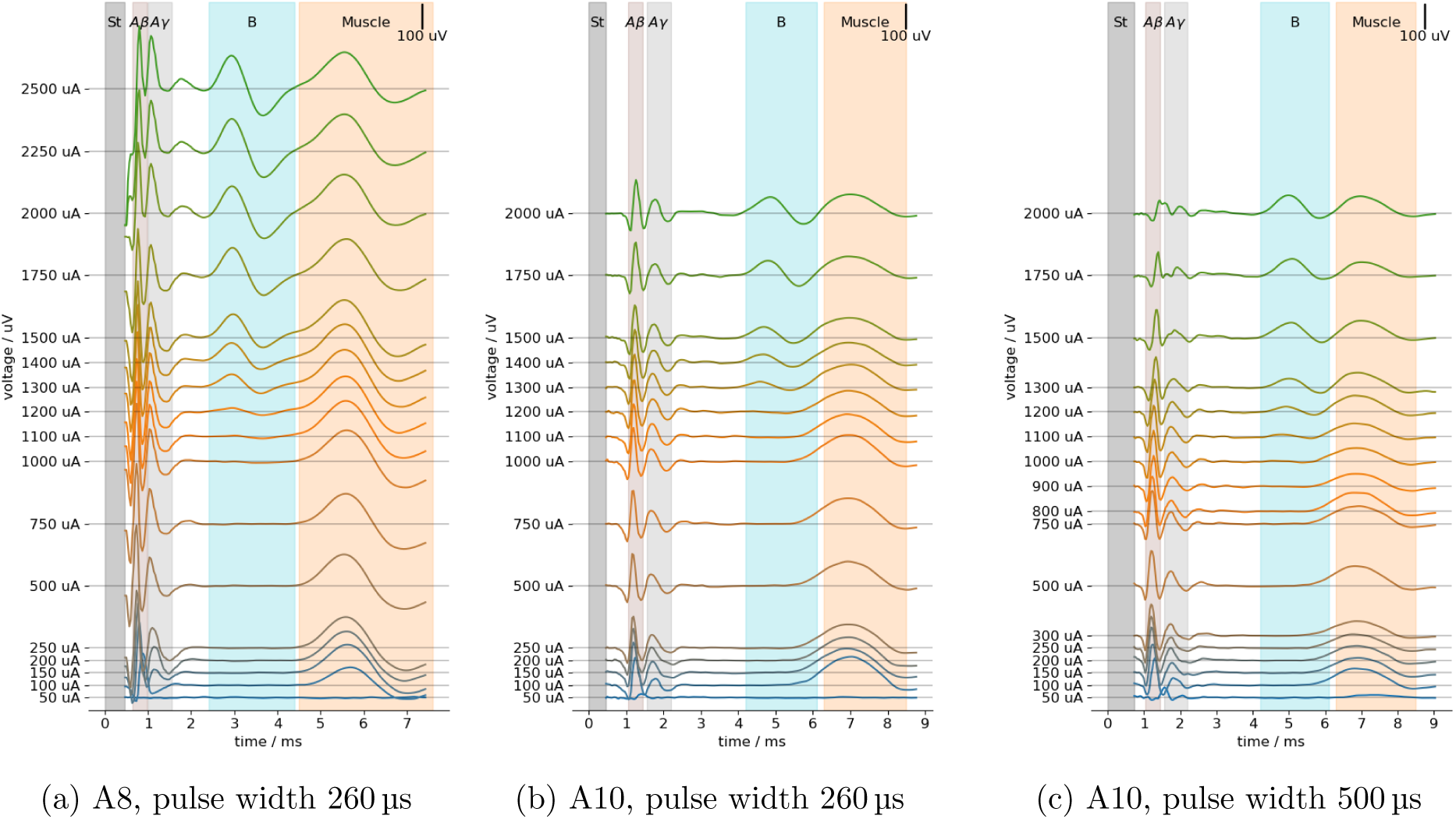
Neural response following stimulation in subjects A8 and A10, Porcine Dataset. The time shown is that from the beginning of the first stimulation pulse (St, the stimulation pulse is not shown). (a) subject A8 at 260 µs. (b), (c) subject A10 at 260 µs and 500 µs. A*β*- and A*γ*-fibre activations emerge at lower stimulation currents than B-fibre activation. Signal artefacts from muscle activation are present in all recordings. The apparent decline in A*β* and A*γ* activation in (c) at high currents is most likely due to interference with a third A-fibre activation between them.

### 3.2 Offline assessment of Bayesian optimization using simulations

Parameter optimization for achieving a specific target response was assessed by applying targeted BO to B-fibre activation as derived from eCAP simulators. One example of a B-fibre activation response is shown in Fig. 6, which is obtained by measuring B-fibre activation in neurograms provided by the eCAPs simulator for A10 (Fig. 5) (similarly for A7, A8, A9 in the supplemental material). A targeted BO was set up to find inputs that yield responses close to a target value of 0.04 mV for A10 (0.06, 0.15, 0.1 mV for A7 to A9). BO was performed with a prior factor of *κ* = 0.8 and and confidence bound factor of *λ* = 2.0. The approximation achieved by surrogate function of the BO after 11 evaluations (including 2 initial input points at the extreme of the search space in magenta) is shown in Fig. 6b. The iso-response curve is indicated by red markers.

**Figure 5:**
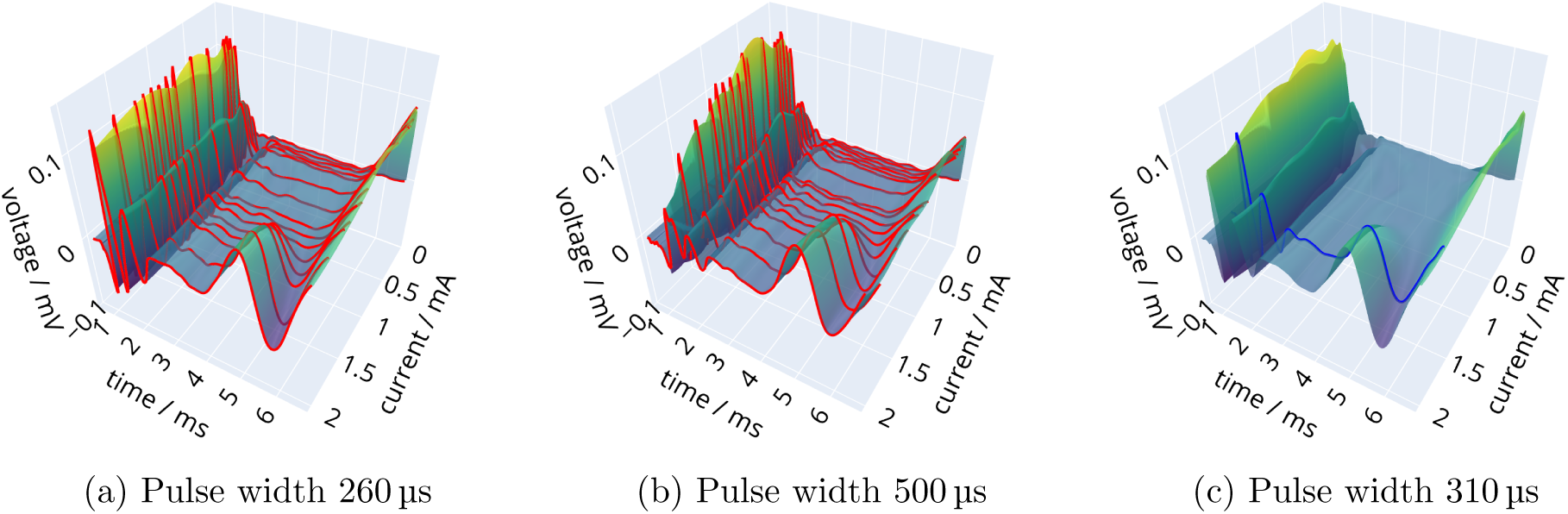
Simulator for neural response to stimulation in A10, Porcine Dataset. The GP simulator is derived from stimulation responses in subject A10 (compare with Fig. 4b and 4c). (a), (b) represent the time response for pulse widths 260 µs and 500 µs. (c) shows a simulated stimulation response surface for pulse width 310 µs and a single stimulation response for pulse current 2.13 mA (blue), which was not observed in the data. The onset of the laryngeal muscle artefact is partly shown.

**Figure 6:**
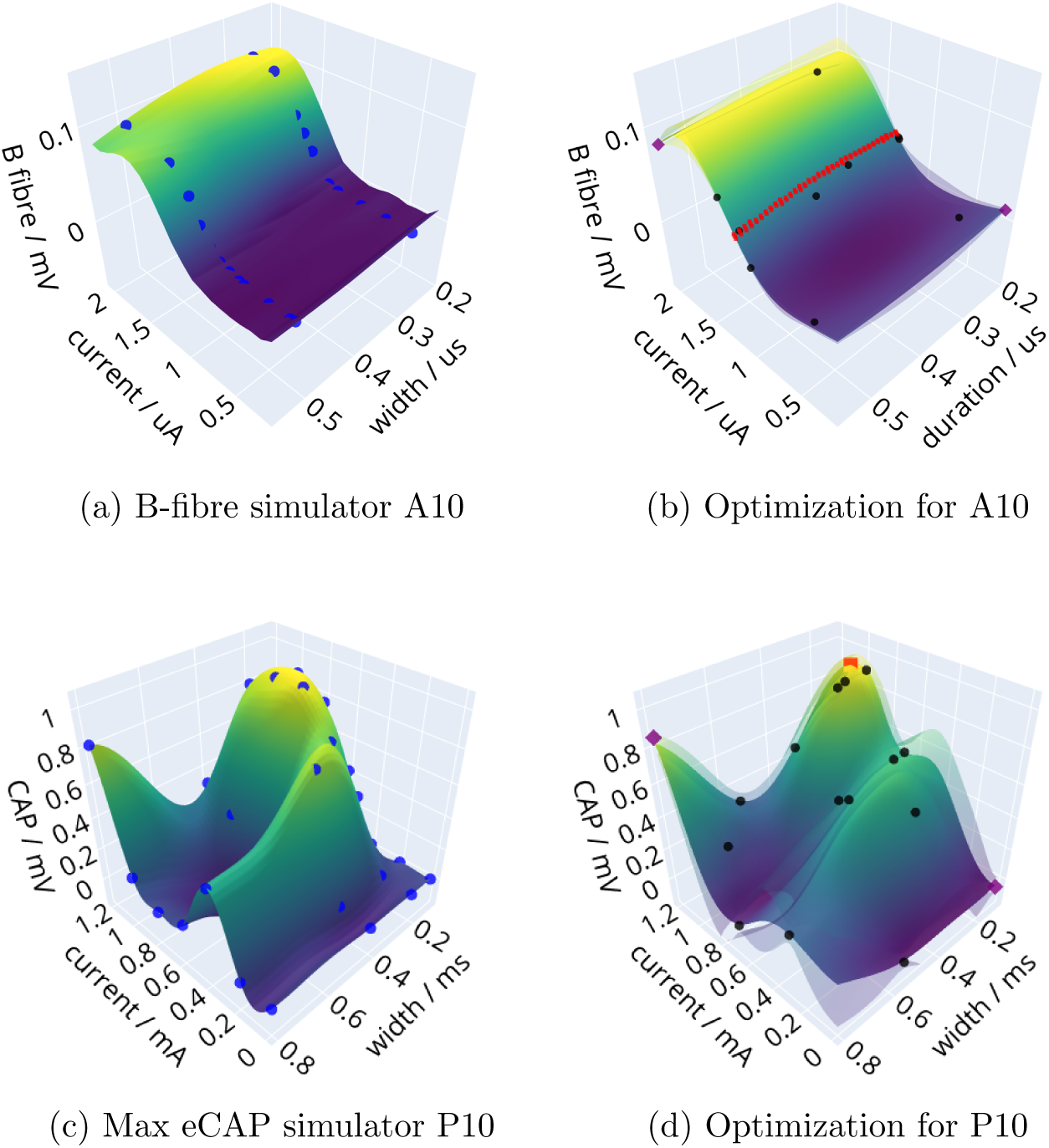
Optimization of B-fibre activation for a simulator for subject A10, Porcine Dataset, and subject P10, Rodent Dataset. (a) B-fibre activation surface derived from neurograms generated by the simulator of Fig. 5, experimental activation values in blue. (b) The surrogate function after a BO series optimizing for a target value of 0.04 requiring 11 BO queries: 2 initial values (magenta), and 9 queries of the BO process. Points of that are closest to the target value along the iso-response curve are shown in red, responses to queries in black. (c) Maximimal eCAP response surface, experimental activation values in blue. (d) The surrogate function after a BO series optimizing for a maximum response (red point) after 15 BO queries: two initial values (magenta), and 13 queries of the BO process. Lower and upper confidence surfaces of the GP surrogate function at one standard deviation are transparent surfaces. BO results for the other subjects can be found in the supplementary material.

Notice how closely the BO’s surrogate approximation in 6b follows the true response in 6a after only a few stimulation results. A small amount of uncertainty is indicated by upper and lower surfaces close to the surrogate approximation.

Similarly, GP simulators were constructed for maximal eCAP activation in 12 subjects, P1 to P12, from the Rodent Dataset. Fig. 6c shows a simulator for subject P10, while Fig. 6d shows the result of optimizing pulse current and width to achieve maximal overall eCAP activation (examples for all 12 subjects can be found in the supplementary material). Each BO was started by two initialisation points (in magenta) at the low and the high value corners of the search spaces. The maximum suggested by the BO is marked by a point in red. Uncertainty estimates around the reconstructed response are indicated by transparent surfaces. They are narrower in the region of interest around the maximum response where BO tends to provide more queries. BO was performed with a prior factor *κ* = 0.8 and confidence bound factor *λ* = 4.0. Again notice the similarity between the BO’s reconstructed surrogate function in 6d compared to the true response in 6c, as well as the BO’s ability to find the the correct maximum for a surface with several local maximima.

In order to evaluate the performance of a BO we compared it to the optimization results from a grid as well as a random search. The results are shown in Fig. 7a for the Porcine and in Fig. 7b for the Rodent Dataset. The BO result (blue markers) is compared to the performance of a random search (grey markers). Also shown are the threshold values provided by the optima in grid searches. The BO matches or outperforms all grid searches after 13 evaluations. The exceptional performance of the grid searches and BO for P3, P4, and P5 in Fig. 7b is explained by the fact that the optima are in or close to one of the corners of the search space and therefore easily captured in a grid search as well as the initial BO evaluation.

**Figure 7:**
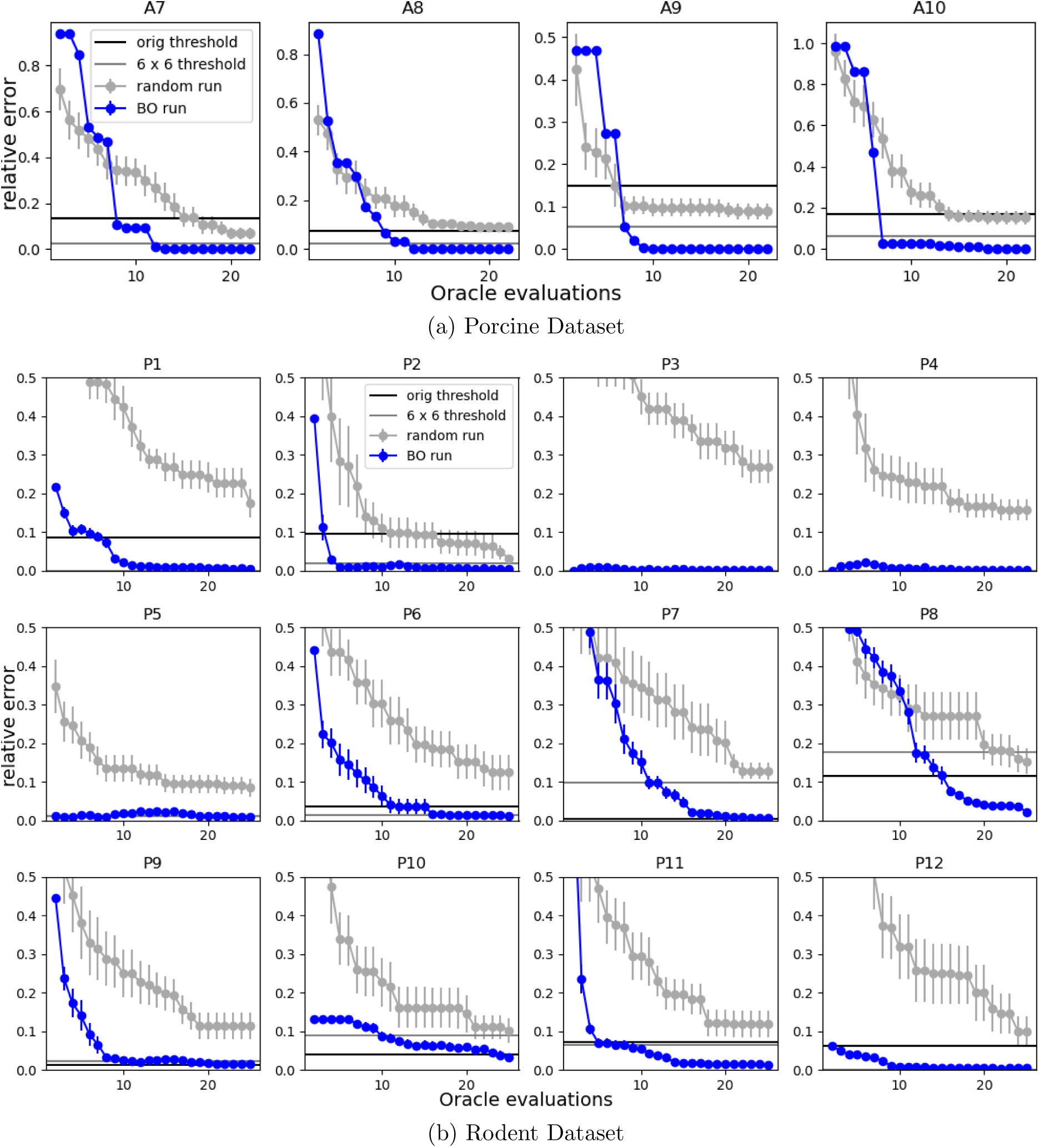
Relative error of BO optimization compared to grid and random optimization. The relative error (blue markers) is relative to the optimal value. For comparison, the results of a random search are shown (grey markers) as mean over 10 runs (with standard error of the mean indicated). Thresholds we expect the BO to reach are provided by optimizing over two grids: the optimum for the original experimental grid (black line) and for a 6 *×* 6 grid (grey line) spanning the input space with 36 points. (a) Evaluation on simulators from subjects A7 to A10, Porcine Dataset (compare with Figs. 6a and 6b for A10). Optimizations were for target values of 0.06, 0.15, 0.1, 0.04 mV for A7 to A10. The experimental data grids were of sizes 33, 34, 20, and 32 for A7 to A10. (b) Evaluation on simulators from subjects P1 to P2, Rodent Dataset. Optimization was for a maximum value. The experimental data grids were all of size 27 (4 *×* 8).

Overall, apart from the greater efficiency in terms of the number of stimulations required, BO also provides a more robust search result compared to grid searches, which are sensitive to the grid layout relative to the optimum input.

### 3.3 Online Bayesian optimization of heart rate change

Simulation results for BO were encouraging and it was decided to apply OBOES online during surgery on porcine subjects to achieve targeted changes in heart rate through manipulation of VNS parameters.

A typical OBOES series with 11 evaluations is shown in Fig. 8 for subject A10 targeting a ΔHR of *−*10 bpm. After two initial evaluations OBOES suggests further VNS parameters. The response to the stimulation is recorded, measured, and forwarded to the algorithm. The progression in updating OBOES’s surrogate function is shown. Notice a narrowing of the upper and lower uncertainty surfaces (transparent) of the GP with a growing number of evaluations. Similar figures for subject A11 can be found in the supplementary material.

**Figure 8:**
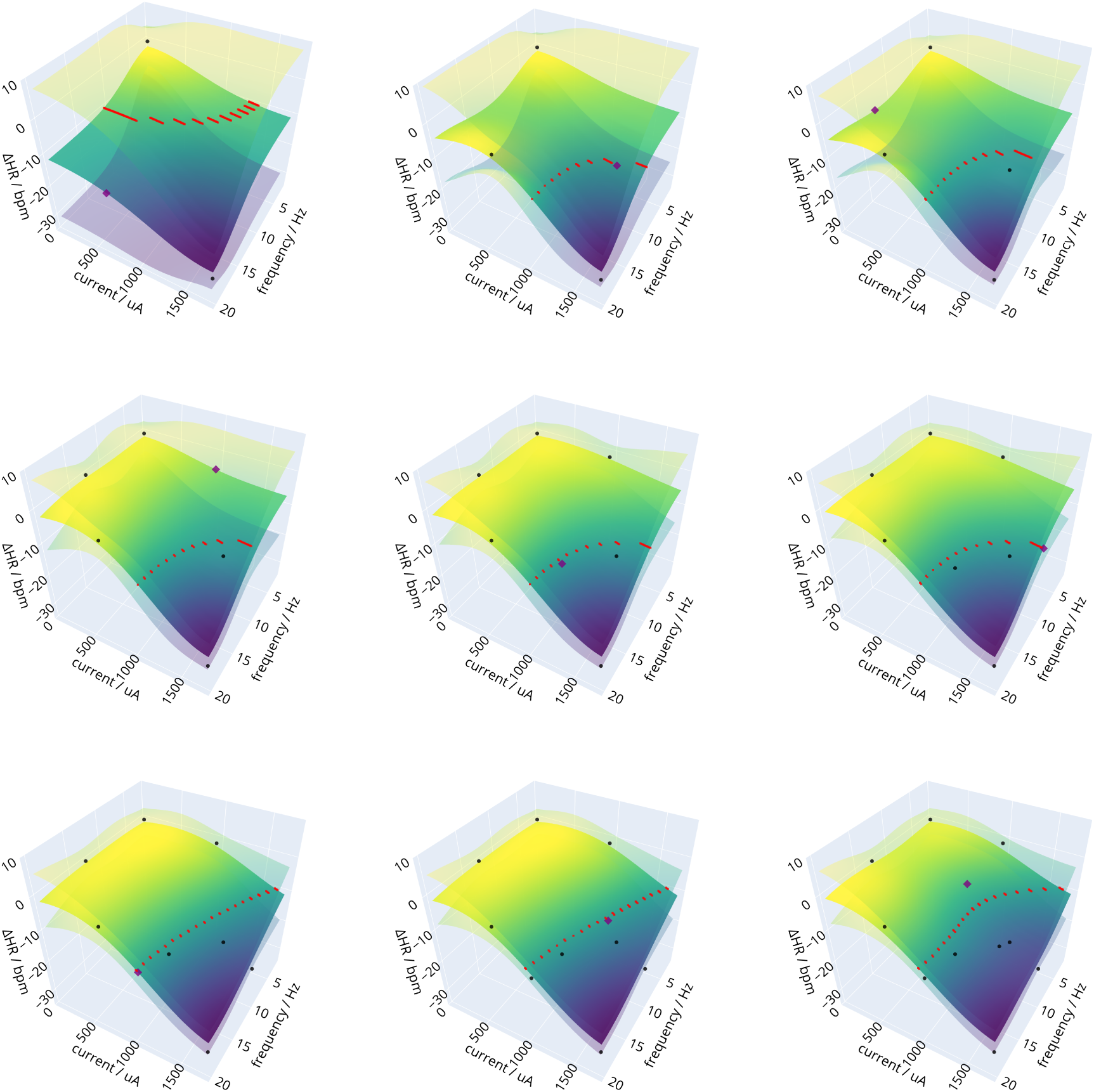
Online OBOES series for subject A10. A series of nine BO steps from two initialization points is shown with a target HR change of -10 bpm. The middle surface represents the mean of the fitted GP, the upper and lower surfaces the upper and lower confidence bound one standard deviation away. Markers shown: observed experimental HR changes (black), the BO query point (magenta) at that stage, potential input points within 0.3 bpm of the target along the iso-response curve (red).

The final result for A10 is shown in Fig. 9 (and for A11 in supplementary material). In order to assess the quality of the surrogate surface three stimulations for each of three different inputs were performed (nine in total) as indicated in Fig. 9a by blue markers. Fig. 9b shows the deviations from the predicted value for each of the three inputs. The posterior probability distribution of the estimated true experimental response is indicated. The expectation is that little probability mass *p* falls outside the GP’s confidence bounds, which is indeed the case (*p <* 0.001). In other words, the GP’s upper and lower confidence surfaces provide reliable guidelines on how much uncertainty stemming from measurement noise and limited knowledge of the true response function affects predicting the effects of optimal stimulation. Since confidence bounds reduce with an increasing number of OBOES steps, they potentially provide a guideline when to stop the optimization procedure, namely, when the prediction uncertainty falls below a specific level of acceptance.

**Figure 9:**
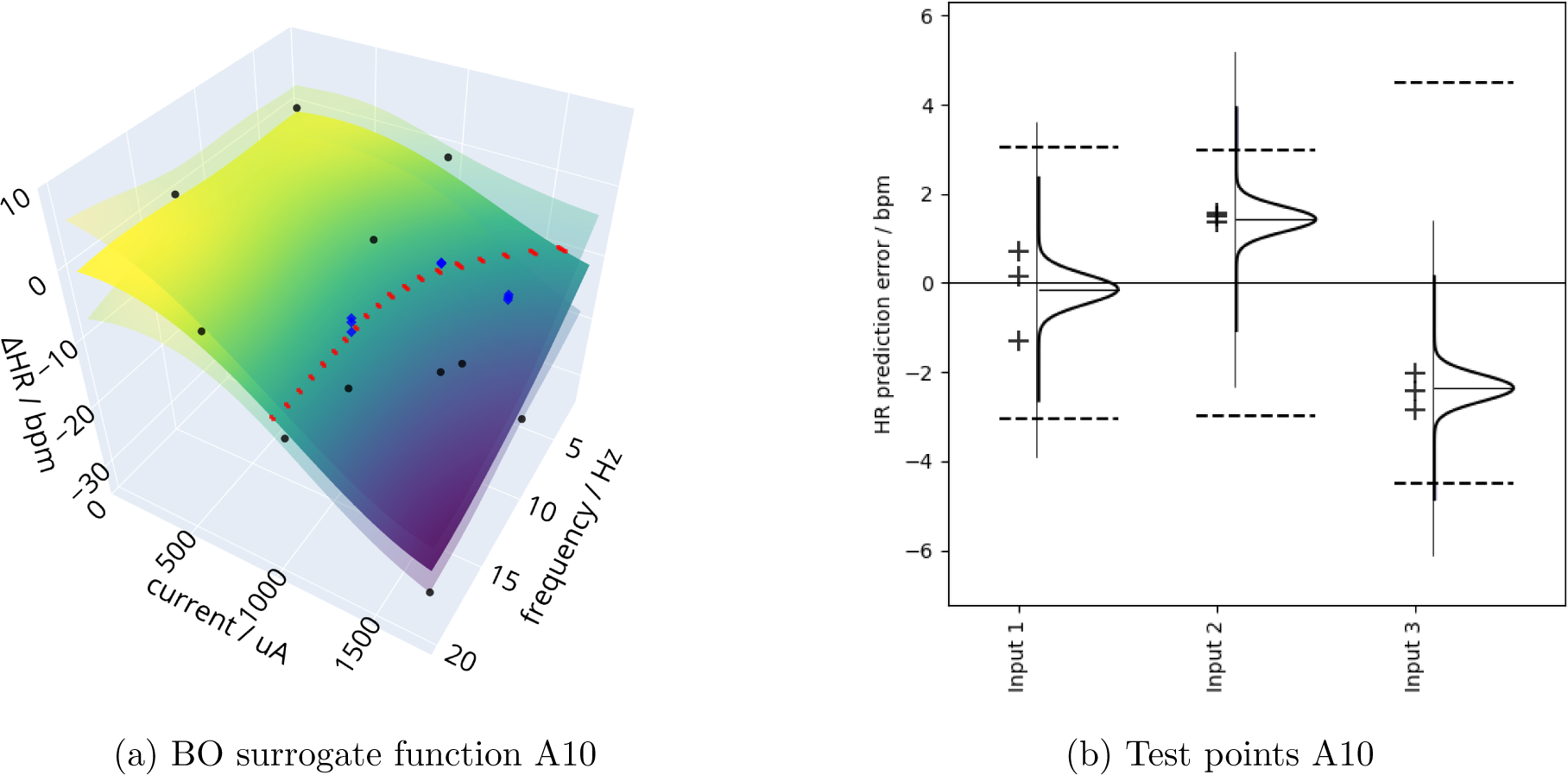
Final test responses after OBOES for target ΔHR for A10. The surrogate models after 11 evaluations for a target HR change of -10 bpm are shown (including two initial points). (a) three responses (blue markers) were measured for each of three different inputs: two selected along the iso-response curve (red), one slightly off target. (b) statistical performance of the test points. The posterior probability distributions of the estimated true HR response based on the three measurements are shown for each evaluation set. The Gaussian process credible interval of one standard deviation represented by transparent upper and lower surfaces in (a) is indicated with broken lines in (b). The probability that the true response falls outside this interval is less than 0.001 in all cases.

### 3.4 Online Bayesian optimization of B fibre activation

The parasympathetic effect of B-fibre activation on heart rate is widely accepted (Qing et al., 2018). Fig. 10 shows a comparison of the effect of stimulation current with that of B-fibre activation on ΔHR in subject A10. This suggests that optimizing B-fibre activation in place of heart rate is a viable alternative to optimizing heart rate directly.

**Figure 10:**
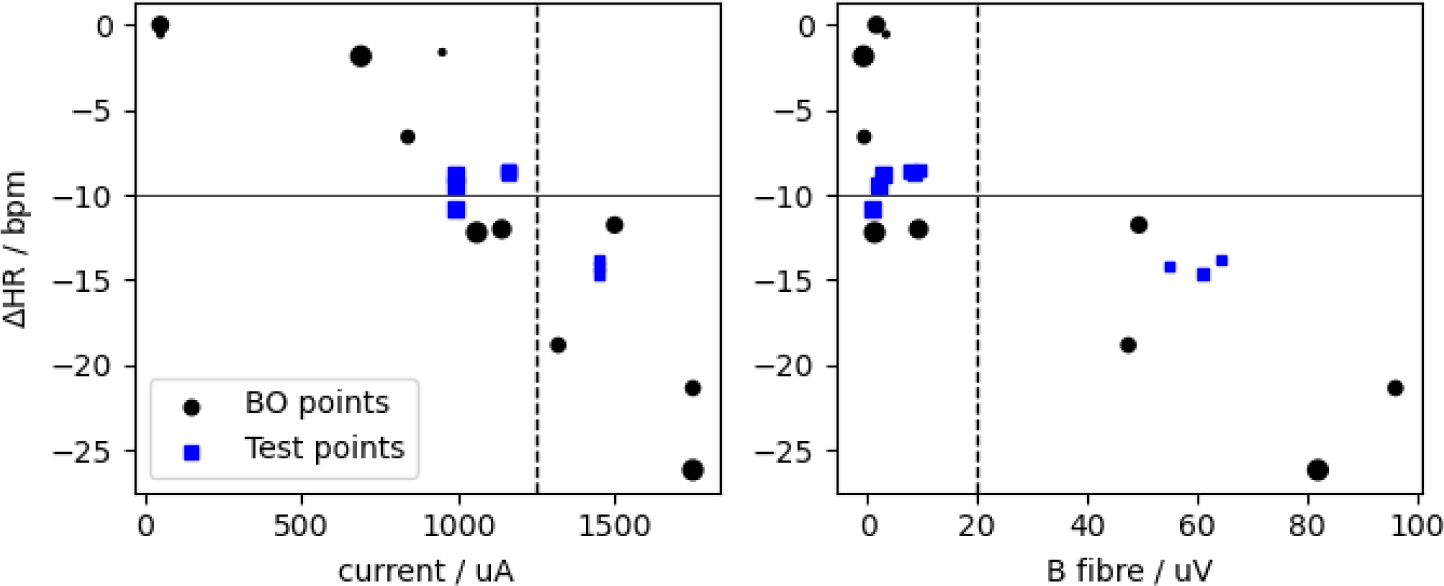
Dependency of ΔHR on pulse current and B fibre activation for stimulations in A10. The frequency is indicated by marker size (smallest 2 Hz, largest 20 Hz). The target ΔHR is indicated by a horizontal line. BO query (black, round) and test stimulations (blue, square) from Fig. 9 show a clear relationship of ΔHR to pulse current (Spearman rank correlation 0.85, *p <* 10*^−^*^5^) and B fibre activation (0.75, *p <* 10*^−^*^4^).

To avoid side effects the total stimulation charge needs to be controlled. In the OBOES experiment of Fig. 11 we aimed to maximize B-fibre activation while minimizing the stimulation current (sequential figures illustrating this optimization in supplementary materials). A way to achieve that is by maximizing the ratio between activation and current. As expected the maximum is achieved around 1500 µA, away from the upper limit that was put on the current by the electrophysiologist. The pulse width was not considered a major cause of potential side effects and was not incorporated as a constraint into the objective function of the BO (i.e. the ratio between B-fibre activation and pulse current). The objective function therefore increases monotonically with the pulse width and the optimum is reached at the maximum of its allowed range, at 500 µs. Fig. 11b demonstrates that the GP confidence range provides a reliable estimate of the uncertainty: the probability *p* that the true response, as estimated from three test points, lies outside the range is negligible (*p <* 0.001). However, although within the uncertainty range, some underestimation bias by the GP of the response is noticeable.

**Figure 11:**
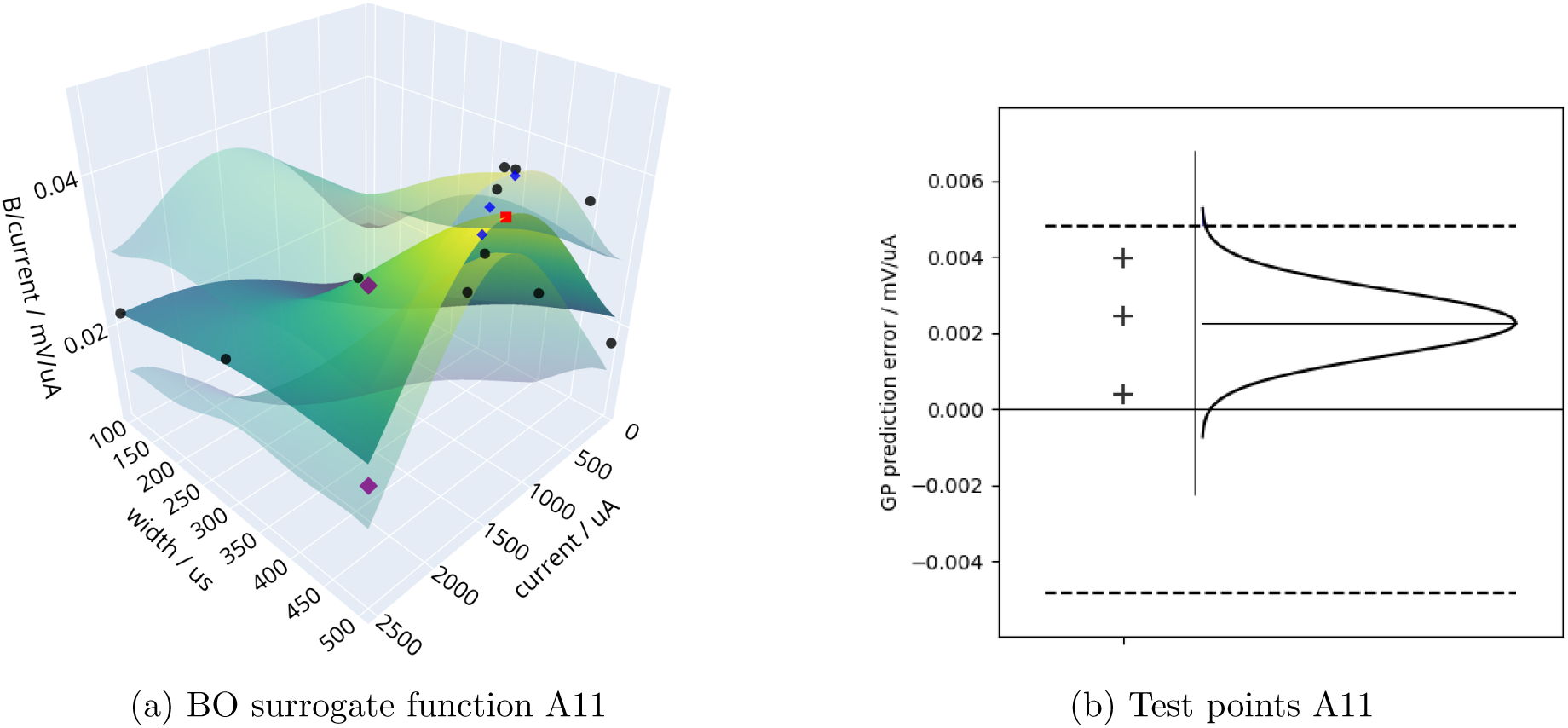
Response surface after a OBOES for maximum B-fibre activation over current for A12. (a) The surrogate model after 13 evaluations with confidence surfaces of 1.5 standard deviations. After OBOES, three test responses (blue markers) were measured around the optimum. (b) Statistical performance of the test points. The posterior distributions of the estimated true activation over current response based on the three test measurements. The Gaussian process credible interval of 1.5 standard deviation represented by transparent surfaces in (a) is indicated with broken lines in (b). The probability that the true response falls outside this interval is less than 0.001.

## 4 Discussion

To our knowledge the OBOES framework presented in this study is the first example of achieving targeted changes in heart rate and maximizing specific fibre activations efficiently through joint optimization of VNS parameters in alive animals. We also present a comprehensive cloud based R&D system enabling real time automated data collection, analysis and optimization. Furthermore, we present a realistic simulation framework modelling responses to VNS. We demonstrate the usage of such simulation models for the development and testing of parameter optimization procedures for VNS.

### 4.1 Performance of OBOES

Current state-of-the-art clinical VNS systems use cardiovascular responses (Ardell et al., 2017b) or tolerance-based dosing strategies (Premchand et al., 2014) to guide VNS parameter choices. This is typically done on a reduced set of parameters, for example, through dose titration using only stimulation current. If multiple parameters are used, they are explored in a rigid order (LivaNova, 2021). The neural stimulation optimization using a Bayesian method, Bayesian optimization as suggested in this study, is more flexible and efficient, particularly for high-dimensional parameter spaces. Bayesian optimization has been successfully used in a variety of other areas of medical research, for example, to guide pharmaceutical dose-optimisation (Takahashi and Suzuki, 2021), or in the design of active neuroprostheses (Losanno et al., 2021). Since the current study we began to apply OBOES to five dimensional parameter spaces including discrete spaces such as electrode locations, which will be reported on in future studies. Due to its speed of optimization, OBOES is better suited to cope with variation in bioelectronic coupling of the electrode-nerve interface at implantation or changes at the interface over time, for example, with inflammation (Qiao, Stieglitz, and Yoshida, 2016).

In the simulation studies, BO performed well compared to grid searches and random searches in terms of the number of steps to get close to an optimum as well as the quality of the approximation (Fig. 7 of section 3.2). The aim of this study, however, was not necessarily to demonstrate superiority over alternative optimization approaches. There are undoubtedly many approaches that perform comparably in terms of requiring only a small number of steps to reach an optimum, BO being one of them. We opted for a BO approach because it (1) enables a principled probabilistic way to integrate prior knowledge, e.g. about the smoothness of the response function, (2) constructs an explicit model of the unknown response function in the form of a surrogate function, and (3) provides information about the amount of model uncertainty. This allows one, for example, to assess the plausibility of the GP’s internal surrogate model and to make a rational assessment of the risk when relying on its predictions in a clinical setting.

A BO approach also enables great flexibility in the choice or design of the acquisition function, which can be adapted to specific requirements. For example, in this study we developed a variant, TCB, of an acquisition function that allows a BO to home in on an iso-response curve of inputs that result in the same defined target value as responses. TCB turned out to be very robust, in simulations as well as in online settings, leading to fast convergence to an iso-response set of inputs. An additional benefit of this approach is that the iso-response curve allows one to pick preferred parameter sets with ancillary benefits, for example, mitigating side effects or promoting device battery life, while maintaining the same biological response.

### 4.2 Online implementations for medical devices

The ITx Platform of Section 2.3 shows two services where a machine learning model can be deployed: either on the acquisition hardware, an edge deployment, or in the ITx Platform portal, a cloud deployment. We opted for the second choice, since we aimed for a proof-of-principle demonstration. A cloud deployment allowed us to manually adjust settings and algorithms and to control stimulus parameters if intervention was deemed necessary. The online BO experiments, however, were surprisingly stable and were obtained with little manual intervention. The use of GP parameter priors stabilised their optimization, particularly in the early phases of the BO where only few data points are available. We have since deployed OBOES directly on the acquisition hardware to run fully automated with the option to monitor and intervene manually if necessary (results not shown here).

Safety considerations are paramount for medical devices, and those leveraging machine learning techniques are no exception. BO is a probabilistic method that balances exploration with optimization of an unknown response function with as few data points as possible. Typically the greatest uncertainty tends to occur at the more extreme values of the input space. A simple method to avoid parameter settings that induce side-effects is to constrain the input parameters to moderate charges initially. As long as the patient’s responses stay within acceptable limits, the size of the input space can be increased gradually, allowing higher charge stimulations to be explored by the model. An alternative approach could exploit the GPs’ natural handling of uncertainty to pursue dose titration in a safe way (Berkenkamp, Krause, and Schoellig, 2016): analysis of the surrogate function can in principle suggest safe extensions of the search space constrained by a risk model that balances expansion of the search space with safety considerations.

### 4.3 Neural biomarkers for physiological effects

Anaesthetics are known to severely dampen the baroreflex and other hemodynamic indices (Ahmed et al., 2021). We observed that to maintain the anaesthetic plane, propofol administration occasionally required adjustment. During grid searches in online experiments, this sometimes drastically changed the magnitude of bradycardia for fixed stimulation parameters. In contrast, nerve recruitment itself is much more robust to anaesthesia or the severity of heart disease. Furthermore, neurograms can be explored rapidly and with few physiological side effects by keeping pulse numbers low (by lowering frequency and train duration). For an intraoperative procedure, optimizing neural biomarkers is thus preferable over optimizing physiological ones. We investigate the relationship between eCAPs and physiological responses in detail in a companion study Berthon et al. (2023). We have shown that OBOES is suitable for optimizing B-fibre activation, a prime example for an indirect optimization of heart rate using a neural biomarker. We also hypothesise that neural biomarkers, based on their enhanced reliability as measures of mechanism engagement and bioelectronic coupling, make good candidates for true closed-loop neuromodulation therapies.

### 4.4 Clinical use of intraoperative optimization

VNS parameter optimization for neural or physiological targets can be performed any time after the implantation of the device. We have successfully performed such optimizations in an awake, freely moving animal (as will be reported in a forthcoming study).

However, intraoperative optimization during clinical implantation, while monitoring fibre engagement, could provide immediate objective feedback on (1) the successful installation of the stimulation lead, (2) whether the intended fibre group is being activated and (3) a range of stimulation parameters (on the iso-response line) that can engage the target mechanism. After implantation, device titration can be guided by the range of stimulation parameters known to generate on-target therapeutic effects, as established during implantation. This could drastically reduce the number of clinical interactions needed following device implantation and the overall time required to achieve optimal postoperative dosing.

## 5 Conclusion

Optimization of VNS parameters for optimizing a target response, particularly in an intraoperative setting, is challenging. We demonstrated that Bayesian optimization is well-suited to support this task. For offline exploration of a wide range of neural biomarkers, accurate simulation models based on Gaussian processes can be derived from experimental data. For online optimization, Bayesian optimization enables an efficient and accurate reconstruction and robust optimization of the response function with comparatively few test stimulations.

## Data availability

A simulation framework for applying OBOES including data used in this study for demonstration purpose is available through the o^2^S^2^PARC platform^1^.

## Authors contribution

LW designed the OBOES algorithms and analyzed the data. LW and TE prepared the first draft of the manuscript. TE, PF-P, OA, MS designed the animal experiments. PF-P and JM guided the animal experiments. LW, AB, OT-L designed, implemented and operated the BO interface of the ITx platform. BP, WB, MJ designed and implemented the cloud based data processing and storage pipeline. BA, WB, SG, MJ designed and implemented the NeuroTool interface. SS, SG, PG, MP, MS, BA, WB, MJ designed and built the BIOS Neural Interface. ES, MT, CH, SL, JJ, AT, GL contributed to reviewing and rewriting of the manuscript. EH, AT, OA, TE conceived of the study. OA supervised the study.

## Footnotes

1 https://osparc.io/study/861f8d70-a2d4-11ed-8c93-02420a0b00f9

